# Neuroinvasive and neurovirulent potential of SARS-CoV-2 in the acute and post-acute phase of intranasally inoculated ferrets

**DOI:** 10.1101/2024.09.21.614276

**Authors:** Feline F. W. Benavides, Edwin J. B. Veldhuis Kroeze, Lonneke Leijten, Katharina S. Schmitz, Peter van Run, Thijs Kuiken, Rory D. de Vries, Lisa Bauer, Debby van Riel

## Abstract

Severe acute respiratory syndrome corona virus 2 (SARS-CoV-2) can cause systemic disease, including neurological complications, even after mild respiratory disease. Previous studies have shown that SARS-CoV-2 infection can induce neurovirulence through microglial activation in the brains of patients and experimentally inoculated animals, which are models representative for moderate to severe respiratory disease. Here, we aimed to investigate the neuroinvasive and neurovirulent potential of SARS-CoV-2 in intranasally inoculated ferrets, a model for subclinical to mild respiratory disease. The presence of viral RNA, histological lesions, virus-infected cells, and the number and surface area of microglia and astrocytes were investigated. Viral RNA was detected in various respiratory tissue samples by qPCR at 7 days post inoculation (dpi). Virus antigen was detected in the nasal turbinates of ferrets sacrificed at 7 dpi and was associated with inflammation. Viral RNA was detected in the brains of ferrets sacrificed 7 dpi, but *in situ* hybridization nor immunohistochemistry did not verify evidence of infection. Histopathological analysis of the brains showed no evidence for an influx of inflammatory cells. Despite this, we observed an increased number of Alzheimer type II astrocytes in the hindbrains of SARS-CoV-2 inoculated ferrets. Additionally, we detected an increased microglial activation in the olfactory bulb and hippocampus, and a decrease in the astrocytic activation status in the white matter and hippocampus of SARS-CoV-2 inoculated ferrets. In conclusion, although showed that SARS-CoV-2 has limited neuroinvasive potential in this model for subclinical to mild respiratory disease, there is evidence for neurovirulent potential. This study highlights the value of this ferret model to study the neuropathogenecity of SARS-CoV-2 and reveals that a mild SARS-CoV-2 infection can affect both microglia and astrocytes in different parts of the brain.

## Introduction

Severe acute respiratory syndrome coronavirus 2 (SARS-CoV-2) infection causes respiratory disease. Additionally, a broad variety of neurological symptoms has been reported occasionally the acute and post-acute phases. These neurological complications can occur after severe, mild or even subclinical respiratory disease and comprise a variety of symptoms. In the acute phase, symptoms including brain fog, headache and memory loss have been observed and several of these symptoms can also persist in the post-acute phase and are part of long-COVID (1,2). Yet, the pathogenesis of neurological disease resulting from SARS-CoV-2 infection is not fully understood.

SARS-CoV-2 has the ability for neuroinvasion, meaning that the virus can enter the peripheral or central nervous system (PNS, CNS) (3). Several studies have shown that SARS-CoV-2 can enter the CNS through axonal transport via several cranial nerves (the olfactory and trigeminal nerves, among others) or via the hematogenous route (3). One important route is via the olfactory mucosa (located in the nasal cavity) where replication of SARS-CoV-2 can result in virus spread through the cribriform plate into the olfactory bulb of the brain. In the olfactory mucosa of coronavirus disease 2019 (COVID-19) patients, SARS-CoV-2 replicates predominantly in sustentacular cells and rarely in olfactory receptor neurons based on the expression of viral antigen or viral RNA (4,5). In tissues obtained from human autopsies, viral RNA or virus antigen was detected in the olfactory bulb (5–8); however, other studies failed to confirm this (8,9). Virus antigen and viral RNA have been detected in neurons, astrocytes and oligodendrocytes in other areas of the human CNS, like the frontal lobe, hypothalamus, basal ganglia, cerebellum and spinal cord (9–13). In experimentally inoculation animals, virus antigen and viral RNA were detected in the olfactory bulb of cynomolgus macaques, ferrets, mice and hamsters (14–18), suggesting neuroinvasion via the olfactory nerve. *In vitro*, SARS-CoV-2 infects neurons in human brain organoids, human pluripotent stem cell (hPSC)-derived dopaminergic neurons and hPSC-derived cortical neurons and astrocytes (18–22). These studies show that SARS-CoV-2 has both a neuroinvasive and neurotropic potential, although the frequency of neuroinvasion is not well understood.

Neurovirulence of SARS-CoV-2 infection is its ability to cause lesions in the CNS leading to clinical disease, independent of direct infection of CNS cells (3). Several mechanisms, such as microgliosis and astrogliosis, are thought to contribute to the neurovirulence of SARS-CoV-2 infection. The activation of microglia (microgliosis) and astrocytes (astrogliosis) is characterized by morphological changes of these cell types as well as their influx into affected areas. In deceased COVID-19 patients, more extensively ramified microglia were detected in the olfactory bulb, white matter and medulla oblongata (11,23,24), as well as an increased influx of microglia into the olfactory bulb, meninges, brainstem and cerebellum (7,11,25–27). Astrogliosis was observed in grey and white matter, in the olfactory bulb, frontal cortex, medulla oblongata, cerebellum and basal ganglia of COVID-19 patients (7,11,24). Transcriptomic analysis comparing post-mortem brain tissues of control and COVID-19 patients revealed a significant enrichment of astrocyte-associated genes in the amygdala potentially indicating activation of astrocytes (6). Similar to humans, microglial activation in the olfactory bulb, hippocampus, cortex and medulla oblongata was also observed in experimentally inoculated mice and hamsters (17,23,24,28). Furthermore, enlargement of astrocytes was observed in the olfactory bulb and hippocampus of hamsters (28), while another study failed to report an influx of astrocytes in the olfactory bulb, cortex, medulla oblongata and hippocampus of SARS-CoV-2 inoculated hamsters (24).

Altogether, *in vivo* studies have shown that SARS-CoV-2 has a neuroinvasive and neurovirulent potential in humans and animal models for moderate to severe respiratory disease, like mouse and hamster. Whether SARS-CoV-2 infection can also induce neurovirulence during a subclinical or mild infection is currently unknown, Therefore we investigated the neuroinvasive and neurovirulent potential of SARS-CoV-2 in the ferret (*Mustela putorius furo*) that develop subclinical to mild respiratory disease. Here, we characterized the neuroinvasive and neurovirulent potential of the ancestral SARS-CoV-2 in intranasally inoculated ferrets, and investigated the presence of histological lesions, viral RNA, and virus-infected cells, as well as numbers and surface areas of microglia and astrocytes in the ferret brains. In this study, ferrets have been examined at the acute and post-acute phase of SARS-CoV-2 infection. We determined acute phase up to 10 dpi, when the virus has not yet been cleared, and the post-acute phase after 10 dpi, which can potentially provide insights into long-COVID.

## Materials and Methods

### Animal experiments and tissues collection

Tissues were included in this study from intranasally inoculated SARS-CoV-2 ferrets from a previous study conducted in compliance with the Dutch legislation (29) for the protection of animals used for scientific purposes (2014, implementing EU Directive 2010/63) and other relevant regulations. All animal experiments were performed under BSL-3 conditions at Erasmus MC. In short, influenza A virus, SARS-CoV-2 and Aleutian Disease Virus seronegative male and female ferrets (Triple F Farms, PA, USA; between 1-2 years old) were intranasally inoculated with 10^5^ TCID_50_/ml SARS-CoV-2 D614G (isolate BetaCoV/Munich/BavPat1/2020). Only female ferrets that did not receive any treatment were included in this study. Four ferrets were sacrificed 7 days post inoculation (dpi), and three ferrets were sacrificed 21 dpi. Three age-matched and sex-matched non-inoculated ferrets were used as negative control for comparison of histological scoring.

The brain, trigeminal nerve, tip of the nose, nasal septum, and the nasal turbinates, which contained olfactory and respiratory mucosa were sampled for histopathological analysis (Fig 1A). Tissues were fixed in 10% formalin for two weeks. Transverse sections of the fixed brain were made at 0.5 cm intervals to evaluate spatial differences in histopathological changes and subsequently embedded in paraffin (Fig 1A).

**Fig 1.**
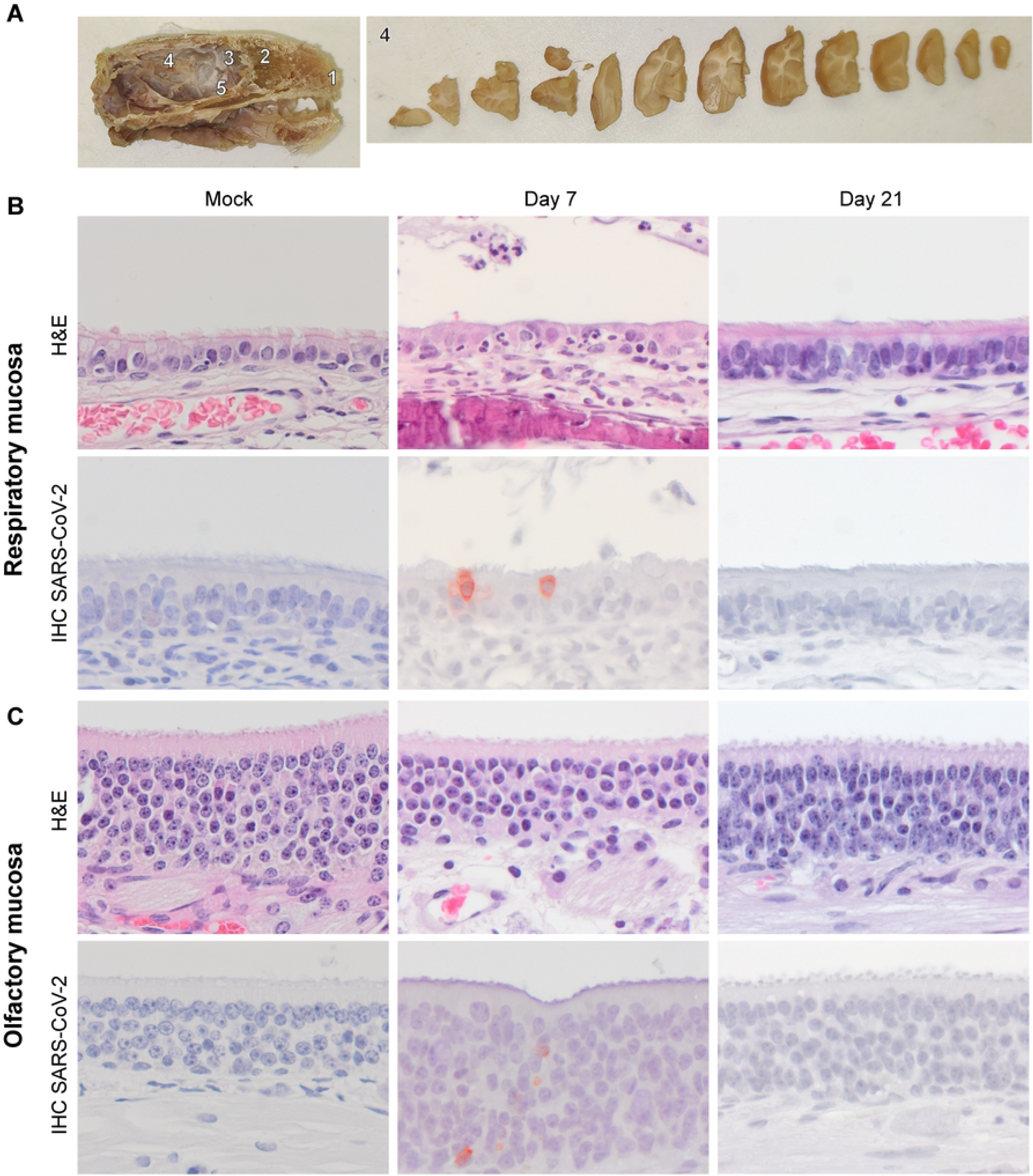
Histology and presence of SARS-CoV-2 antigen expression in the respiratory and olfactory mucosae of SARS-CoV-2 inoculated ferrets. Intranasally inoculated ferrets (D614G; 10^5^ TCID_50_) were sacrificed at day 7 or day 21 post inoculation. Tissues sampled are depicted in (A) left panel: 1 = tip of the nose; 2 = nasal cavity containing respiratory and olfactory mucosa; 3 = olfactory bulb; and 4 = the brain within the cranial cavity; 5 = trigeminal nerve. As shown in the right panel, transverse sections of the fixed brain were made at 0.5 cm intervals to evaluate spatial differences in histopathological changes. Hematoxylin and eosin (H&E) staining and immunohistochemistry (IHC) staining for SARS-CoV-2 nucleoprotein of the respiratory mucosa (B) and olfactory mucosa (C) of SARS-CoV-2 inoculated ferrets. Individual columnar ciliated epithelial cells in the respiratory mucosa collected at 7 dpi express cytoplasmic SARS-CoV-2 antigen (B, lower row) colocalized with infiltration of neutrophils, and intraluminal presence of cellular debris also containing few neutrophils (B, upper row).

### Immunohistochemistry and histopathology

Three-µm-thick formalin-fixed paraffin-embedded (FFPE) tissue sections were deparaffinized, rehydrated and stained with haematoxylin (Klinipath) and eosin (QPath; H&E) to assess histopathological changes. For immunohistochemistry (IHC), consecutive tissue sections were deparaffinized, rehydrated and pre-treated for antigen retrieval. Pre-treatment of slides consisted of boiling for 15 minutes in citric acid buffer (pH 6.0) for the detection of SARS-CoV-2 nucleoprotein (NP) and GFAP (glial fibrillary acidic protein; astrocytes); or boiling for 15 minutes in TRIS-EDTA buffer (pH 9.0) for the detection of IBA1 (ionized calcium binding adaptor molecule 1; microglia). After antigen retrieval, slides were incubated with 3% hydrogen peroxidase (Sigma-Aldrich) for 10 minutes to block endogenous peroxidases. After blocking, slides were washed with phosphate-buffered saline (PBS), and PBS/0.05% Tween 20 (Merck). Sections for SARS-CoV-2 IHC were additionally blocked with 10% goat serum (X0907, DAKO) for 30 minutes at room temperature (RT). Slides were incubated with a rabbit polyclonal antibody against SARS-CoV/SARS-CoV-2-nucleoprotein (40143-T62, Sino Biological, 1:1000), rabbit anti-IBA1 (019-19741, Wako Pure Chemical Corporation; 2.5 µg/ml) or mouse anti-GFAP (clone 4A11, BD556327, Biosciences, 20 µg/ml) in PBS/0,1% BSA (Aurion) for 1h at RT. For each staining on each tissue an isotype control was included as a negative control, either a rabbit IgG isotype control (AB-105-C, R&D, 1:200; 2.5 µg/ml), or mouse IgG2b isotype control (MAB0041, R&D, 5 µg/ml). Tissue sections from negative control ferrets were included as negative tissue controls for all stainings. A lung section from an experimentally inoculated hamster with SARS-CoV-2 was used as positive control for the NP SARS-CoV-2 IHC. After washing, sections were incubated with peroxidase labelled goat-anti-Rabbit IgG (P0448, DAKO, 1:100) or goat-anti-mouse IgG (PO447, Southern Biotech, 1:100) in PBS/0,1% BSA for 1 hour at RT. Peroxidase activity was revealed by incubating slides in 3-amino9-ethylcarbazole (AEC; Tokyo Chemical Industry) in N,N-dimethylformamide (DMF; Sigma-Aldrich) for 10 minutes, resulting in bright red precipitate, followed by counterstaining with haematoxylin (Sigma-Aldrich).

### RNA isolation and qPCR

RNA was extracted from paraffin sections of all available tissues using the RNeasy FFPE kit (Qiagen) according to the manufacturer’s protocol. In short, sections of 5 µM were collected in Eppendorf vials and deparaffinized using xylene. The sample was lysed with proteinase K digestion followed by heat and DNAse treatment. Ethanol and RBC buffer from the kit was added to bind total RNA to the spin columns. Samples were washed and eluted twice in RNAse free water. qPCR was performed targeting E-gene of SARS-CoV-2 as previously reported (30). For the qPCR, the maximum RNA input was used, which corresponds to 100 ng input. The housekeeping gene hypoxanthine phosphoribosyltransferase 1 (HPRT) was included as control; its sequence can be found in (31). All reactions were run on a 7500 Real Time PCR Cycler (Applied Biosystems) in technical duplicates. A Ct-value < 45 was considered positive. Values are depicted ± SD. For the brain, all separate tissue sections were tested individually, but the average is depicted ± SD.

### SARS-CoV-2 in situ hybridization

BaseScope™ RNA probes were designed by Bio-Techne Ltd (Abingdon, UK) for SARS-CoV-2 BA-V-CoV-Wuhan-Nucleocapsid-3zz-st (846661). In situ hybridization (ISH) was performed on FFPE consecutive sections using BaseScope™ Reagent Kit v2–RED (323900) as described by the manufacturer. SARS-CoV-2 nucleocapsid RNA molecules were visualized as red chromogenic dots.

### Microscopy

H&E sections of all tissues were evaluated for histopathological changes with an Olympus BX51 light microscope by an certified veterinary pathologist. A 40× digital scan was made of H&E slides of the hindbrain (area surrounding the pons and cerebellum) for counting Alzheimer type II astrocytes using a Hamamatsu Nanozoomer 2.0 HT digital slide scanner with the accompanying software (NanoZoomer Digital Pathology and NDP.scan and NDP.view, Hamamatsu, Higashiku, Hamamatsu City, Japan). For microglial and astrocytic evaluation, a 10× air objective (Olympus) was used to select a region of interest with an Olympus BX51 microscope. The regions of interest for microglial and astrocytic evaluation in this study were determined as the glomerular layer and the granular cell layer of the olfactory bulb, the white matter tract (corpus callosum), grey matter (posterior parietal cortex) and the hippocampus (CA1 region). We chose these areas, like the corpus callosum, to consistently sample in the same brain area, and therefore a ferret brain atlas was used to determine the correct area for imaging (32). The slides were blinded before the images of the selected regions of interest were taken. Images were taken at either a 200× magnification (20× air objective; Olympus) or 400× magnification (40× air objective; Olympus) with CellSense software.

### Quantitative analysis of IBA1^+^ cells, GFAP^+^ cells and Alzheimer type II astrocytes

At least two sections per brain area were selected per ferret, and per section at least three images were taken of the region of interest in the brain. The number of IBA1^+^ (microglia) and GFAP^+^ (astrocytes) cells per high power field (400× magnification) was determined by manual counting by two observers blinded to the infection status. The averages ± SD of the IBA1^+^ or GFAP^+^ cells per ferret were plotted. As a measure for activation of astrocytes and microglia, the surface areas of IBA1^+^ and GFAP^+^ cells was determined using the pixel classifier function in QuPath 0.4.4 (33) with a Gaussian filter. The threshold values to determine the positive and negative surface areas can be found in Table 1. The averages ± SD of the IBA1^+^ and GFAP^+^ cells surface areas per ferret were plotted. Additionally, the surface area per IBA1^+^ or GFAP^+^ cell was plotted ± SD.

**Table 1.** The chosen threshold values used in QuPath to determine the IBA1^+^ and GFAP^+^ surface area.

| Region of interest | IBA1<sup>+</sup> surface area | GFAP<sup>+</sup> surface area |
| --- | --- | --- |
| Glomerular layer (olfactory bulb) | 0.12 | N.P. |
| Granular cell layer (olfactory bulb) | 0.12 | N.P. |
| White matter (corpus callosum) | 0.23 | 0.20 |
| Grey matter (posterior parietal cortex) | 0.18 | 0.21 |
| Hippocampus (CA1 region) | 0.17 | 0.17 |

N.P. = not performed, IBA1 = ionized calcium binding adaptor molecule 1, GFAP = glial fibrillary acidic protein.

Alzheimer type II astrocytes were counted manually from a 40× digital scan of a H&E slide in Qupath 0.4.4 if the cell matched to the description referenced before (34,35). Alzheimer type II astrocytes are characterized by a large, pale nucleus, with a rim of chromatin and a small conspicuous nucleolus. The total brain area in the slide was calculated in µm^2^ in Qupath 0.4.4 (33), and the number of Alzheimer type II astrocytes was normalized to number of cells per mm^2^ brain area and plotted ± SD.

### Statistical analysis

Statistical differences between experimental groups were determined by using a one-way analysis of variance (ANOVA) with a Dunnett’s *posthoc* test using GraphPad Prism version 10.1.2. P values of ≤ 0.05 were considered statistically significant. Figures were prepared with Adobe Photoshop 22.1.1 and Adobe Illustrator 25.1.0.

## Results

### Assessment of ferret respiratory tissues

To evaluate cell tropism in the respiratory tract of D614G SARS-CoV-2 intranasally inoculated ferrets were sacrificed at 7 or 21 days post inoculation (dpi) and samples of the tip of the nose, nasal septum, nasal turbinates containing olfactory and respiratory mucosa were analysed. As reported earlier, SARS-CoV-2 RNA was detected by qPCR in swabs of the throat and nose of these ferrets still at 7 dpi (29). Additionally, infectious virus was detected in throat swabs of these ferrets at 3 dpi (29). In our analyses on the FFPE tissues, at 7 dpi, viral RNA was detected in tissues of the nasal septum and nasal turbinates in all ferrets (Table 2). At 21 dpi, viral RNA was detected in the tip on the nose (1 out of 3 ferrets) and nasal turbinates (2 out of 3 ferrets; Table 2). ISH verified the presence of the SARS-CoV-2 viral RNA in all qPCR-positive respiratory tissue sections (Table 2). Next, to determine the location and cellular tropism of SARS-CoV-2 in the ferret respiratory tract, IHC for SARS-CoV-2 NP was performed. Virus antigen could not be detected in the tip of the nose or nasal septum from ferrets sacrificed at 7 and 21 dpi. Virus antigen was detected at 7 dpi in both respiratory and olfactory mucosa of the nasal turbinates, but was more pronounced in the ciliated respiratory mucosa cells (Fig 1B). The presence of virus antigen was associated with inflammation, characterized by slight to mild infiltration of neutrophils within the mucosal lamina propria and within the epithelial lining in the respiratory mucosa and in a lesser extent in the olfactory mucosa. Additionally, necrotic sloughed epithelial debris admixed with few neutrophils was present within nasal passages along the inflamed mucosae in one ferret (Fig 1B). Virus antigen or histopathological changes of significance were not detected in the respiratory tissues of ferrets sacrificed at 21 dpi, although sporadic numbers of neutrophils were still present within the lamina propria and within the epithelial lining of the respiratory mucosa of two ferrets (Fig 1C).

**Table 2.** Ct-values of viral RNA detected in various tissues of SARS-CoV-2 inoculated ferrets.

|  |  |  | Ferret |  |  |  |  |  |  |
| --- | --- | --- | --- | --- | --- | --- | --- | --- | --- |
|  | Days post inoculation | Target | #1 | #2 | #3 | #4 | #5 | #6 | #7 |
| Nasal turbinates | 7 | S2 | 32.2 | 28.3 | 27.0 | 32.8 |  |  |  |
|  |  | HPRT | 37.9 | 37.2 | 34.5 | 34.1 |  |  |  |
|  | 21 | S2 |  |  |  |  | 37.3 | N.D. | 32.8 |
|  |  | HPRT |  |  |  |  | 39.0 | 34.9 | 34.8 |
| Tip of the nose | 7 | S2 | N.D. | N.D. | N.D. | N.D. |  |  |  |
|  |  | HPRT | 36.1 | 35.5 | 37.2 | 34.7 |  |  |  |
|  | 21 | S2 |  |  |  |  | N.D. | 39.1 | N.A. |
|  |  | HPRT |  |  |  |  | 36.8 | 36.4 | N.A. |
| Nasal Septum | 7 | S2 | 36.6 | 35.9 | 34.1 | 33.0 |  |  |  |
|  |  | HPRT | 35.9 | 38.2 | 37.2 | 34.2 |  |  |  |
|  | 21 | S2 |  |  |  |  | N.D. | N.D. | N.D. |
|  |  | HPRT |  |  |  |  | 39.5 | 35.8 | 38.9 |
| Trigeminal | 7 | S2 | N.D. | N.D. | N.D. | N.A. |  |  |  |

| nerve |  | HPRT | 37.2 | 37.7 | 37.1 | N.A. |  |  |  |
| --- | --- | --- | --- | --- | --- | --- | --- | --- | --- |
|  | 21 | S2 |  |  |  |  | N.D. | N.A. | N.D. |
|  |  | HPRT |  |  |  |  | 38.7 | N.A. | 42.0 |
| Brain<br>(pooled) | 7 | S2 | 41.5 | 39.0 | 38.6 | 38.5 |  |  |  |
|  |  | HPRT | 37.5 | 36.6 | 35.4 | 35.3 |  |  |  |
|  | 21 | S2 |  |  |  |  | N.D. | N.D. | N.D. |
|  |  | HPRT |  |  |  |  | 37.1 | 35.4 | 36.1 |

S2 = SARS-CoV-2; HPRT = hypoxanthine phosphoribosyltransferase 1 (housekeeping gene); N.D. = not detected, Ct-value $\geq 45$ ; N.A. = not available; Nasal turbinates contain both respiratory and olfactory mucosa; Brain (pooled) is pooled data from all positive qPCR brain tissue sections.

### Assessment of the neuroinvasive potential of SARS-CoV-2 in ferrets

Next, we investigated the neuroinvasive potential of D614G SARS-CoV-2 in ferrets, by testing for viral RNA by qPCR and virus antigen expression by IHC. If viral RNA was detected, we verified this by ISH. Viral RNA was detected by qPCR in inoculated ferrets sacrificed at 7 dpi in various parts of the brain (pooled results of various qPCR positive brain tissues sections in Table 2). However, virus antigen was not detected by IHC in any ferrets and ISH did not confirm the presence of viral RNA in qPCR-positive brain tissues.

### Assessment of histopathological changes in the brain

All brain areas were screened for histopathological abnormalities. We did not detect infiltration of any inflammatory cells, and no gliosis or necrosis in the brain tissues of any ferret. Alzheimer type II astrocytes were detected in the area of the pons and the cerebellum (example in Fig 2A). We observed significantly more Alzheimer type II astrocytes in SARS-CoV-2 inoculated ferrets sacrificed at 7 and 21 dpi compared to control (Fig 2B). The number of Alzheimer type II astrocytes in ferrets sacrificed at 21 dpi were significantly lower compared to ferrets sacrificed at 7 dpi (Fig 2B).

**Fig 2.**
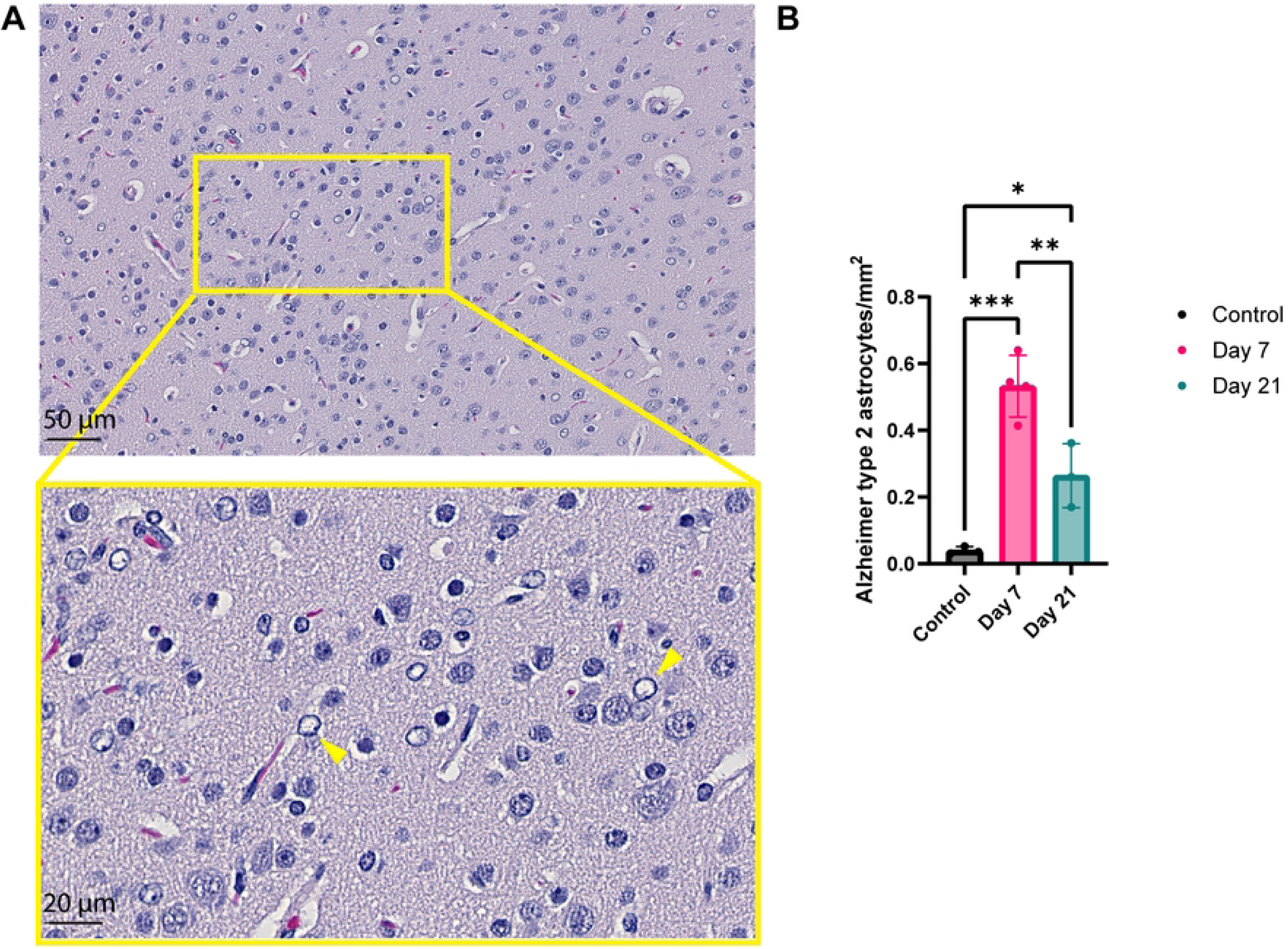
Detection of Alzheimer type II astrocytes in the area of the pons and cerebellum of inoculated ferrets. (A) A 40× digital scan was used for counting Alzheimer type II astrocytes (example with yellow arrows), which were characterized by having a large, pale nucleus with a rim of marginated chromatin and small conspicuous nucleolus. (B) The number of Alzheimer type II astrocytes were counted in a brain slice and normalized to number of cells per mm^2^. Statistical significance was calculated with a one-way analysis of variance (ANOVA) with a Dunnett’s *posthoc* test. Averaged values of three or four individual ferrets per infection group were compared to values of three control ferrets. Data is shown as mean ± SD. Asterisks indicate statistical significance (*P<0.05, **P<0.01, ***P<0.001).

### Assessment of neurovirulence in the olfactory bulb

Expression of IBA1 (microglia) and GFAP (astrocytes) was determined in the brain of D614G inoculated ferrets by IHC to characterize neurovirulence. IBA1 expression in the glomerular layer of olfactory bulbs appeared to be modestly increased by qualitative visual examination in ferrets sacrificed at 7 dpi and 21 dpi compared to the control ferrets (Fig 3A). Quantitative analysis revealed no significant differences in the number of IBA1^+^ cells between control and infected ferrets in both the glomerular and granular cell layers of the olfactory bulb (Fig 3B, C). However, there was a significant increase in the IBA1^+^ surface area per high power field (HPF) in the glomerular layer of the olfactory bulb in ferrets sacrificed at 7 and 21 dpi compared to control ferrets (Fig 3B). Analysis of the surface area normalized per IBA1^+^ cell did not differ significantly between infected and control ferrets in the glomerular layer and granular cell layer of the olfactory bulb (Fig 3B, C). The GFAP expression did not appear to be different between control and infected ferrets (Fig 3D). Quantification of the number and surface area of GFAP^+^ cells was not feasible in the olfactory bulb, because GFAP expression was to widespread and therefore it was not possible to distinguish single cells, which is required for quantification.

**Fig 3.**
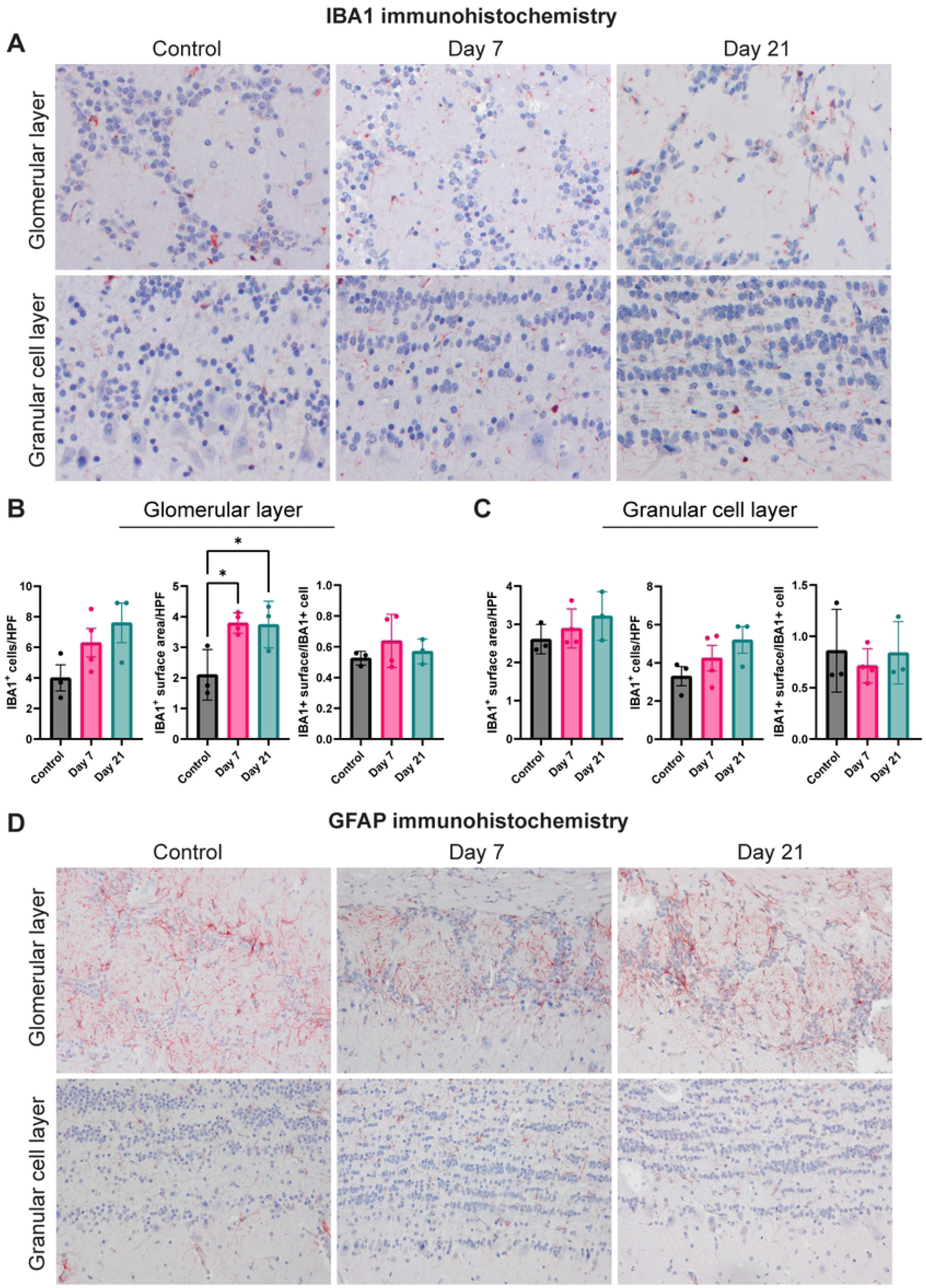
Activation status of microglia and astrocytes in the olfactory bulb of SARS-CoV-2 inoculated ferrets. (A) Detection of microglial (IBA1^+^) cells in the glomerular and granular cell layer of the olfactory bulbs of SARS-CoV-2 inoculated ferrets. The number of IBA1^+^ cells and IBA1^+^ surface area in the glomerular layer (B) and granular cell layer (C) of the olfactory bulb. Statistical significance was calculated with a one-way analysis of variance (ANOVA) with a Dunnett’s *posthoc* test. Averaged values of three or four individual ferrets per infection group were compared to values of three control ferrets. Data shown as mean ± SD. Asterisks indicate statistical significance (*P<0.05). (D) Detection of astrocytes (GFAP^+^) cells in the glomerular and granular cell layer of olfactory bulb of inoculated ferrets. Abbreviations: HPF = high power field, IBA1 = ionized calcium binding adaptor molecule 1, GFAP = glial fibrillary acidic protein.

### Assessment of neurovirulence in, white and grey matter of cerebral cortex and the hippocampus

IBA1 and GFAP IHC was performed to investigate the activation status of microglia and astrocytes, respectively, in the white and grey matter of the cerebral cortex (corpus callosum and posterior parietal cortex, respectively) and the hippocampus (CA1 region). Again, quantitative analyses were performed per brain area: counting the number of IBA1^+^ or GFAP^+^ cells and measuring the IBA1^+^ or GFAP^+^ surface area per HPF (Fig 4). The main trends were as follows. In the white matter, no statistically significant differences in the number of IBA1^+^ cells, surface area or surface area per IBA1^+^ cell were observed (Fig 4A), while significantly less GFAP^+^ surface area was detected (Fig 4B). In the grey matter, no statistically significant differences were observed in the number of IBA1^+^ and GFAP^+^ and their surface area (Fig 4C, D). In the hippocampus, a statistically significant increase in IBA1^+^ surface area was detected (Fig 4E) in inoculated ferrets sacrificed at 7 and 21 dpi compared to control ferrets, while a decrease in GFAP^+^ surface was seen in ferrets sacrificed at 7 dpi compared to control ferrets (Fig 4F).

**Fig 4.**
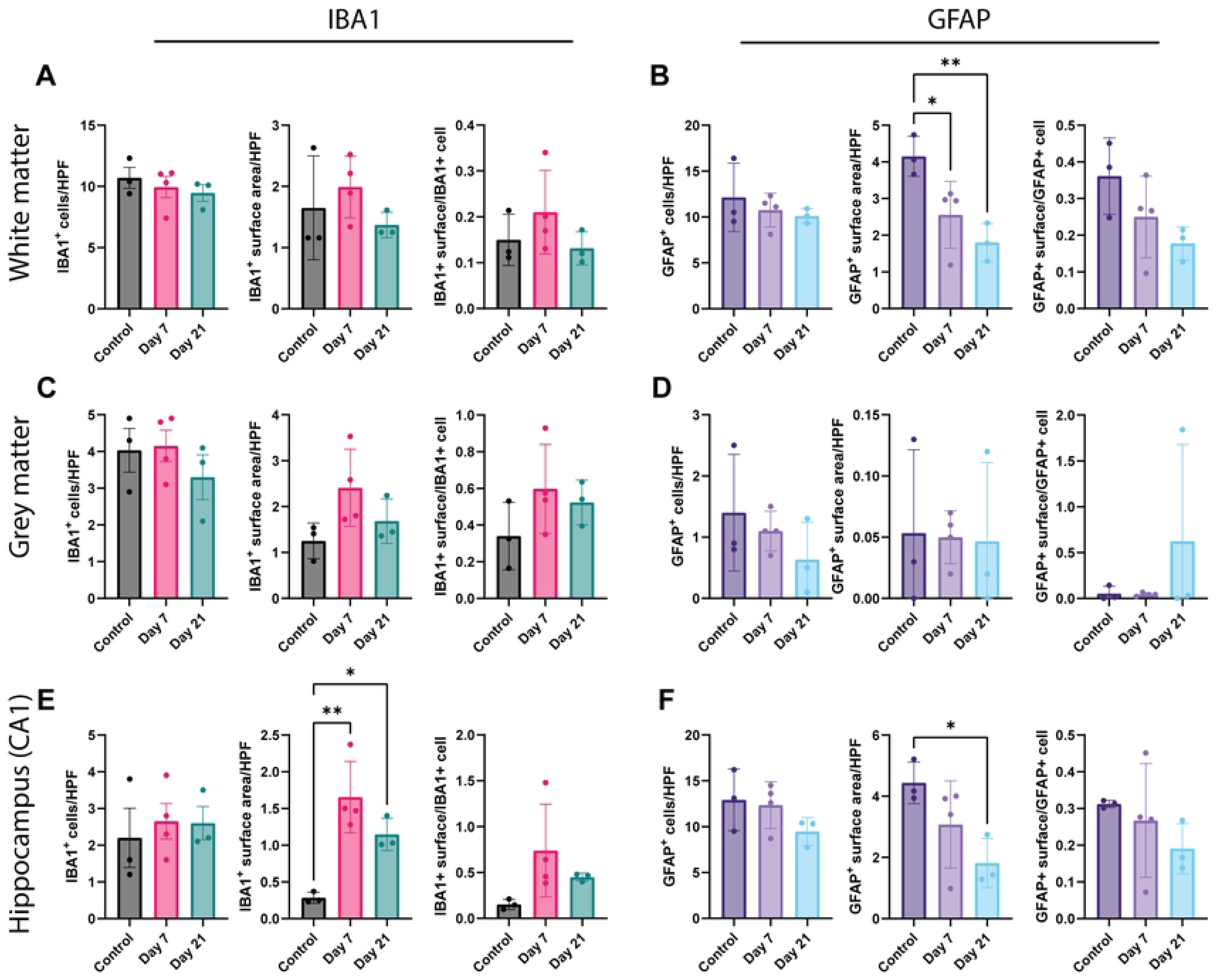
The activation status of microglia and astrocytes in the white matter, grey matter and hippocampus of SARS-CoV-2 infected ferrets. (A) Number of IBA1^+^ cells, the IBA1^+^ surface area and the surface area per cell in the white matter. (B) Number of GFAP^+^ cells, the GFAP^+^ surface area and the surface area per cell in the white matter. (C) Number of IBA1^+^ cells, the IBA1^+^ surface area and the surface area per cell in the grey matter. (D) Number of GFAP^+^ cells, the GFAP^+^ surface area and the surface area per cell in the grey matter. (E) Number of IBA1^+^ cells, the IBA1^+^ surface area and the surface area per cell in the hippocampus (CA1 region). (F) Number of GFAP^+^ cells, the GFAP^+^ surface area and the surface area per cell in the hippocampus (CA1 region). Statistical significance was calculated with a one-way analysis of variance (ANOVA) with a Dunnett’s *posthoc* test. Averaged values of three or four individual ferrets per infection group were compared to values of three control ferrets. Data shown as mean ± SD. Asterisks indicate statistical significance (*P<0.05, **P<0.01). Abbreviations: HPF = high power field, IBA1 = ionized calcium binding adaptor molecule 1, GFAP = glial fibrillary acidic protein.

## Discussion

This study demonstrates that SARS-CoV-2 has limited neuroinvasive potential via the olfactory route in this ferret model, although neuroinvasion via the hematogenous route or neuroinvasion at earlier timepoints cannot be excluded. We observed SARS-CoV-2 replication in the respiratory and olfactory mucosa confirming other studies (4,5,14,16,17,29). The ability of SARS-CoV-2 to replicate in the olfactory mucosa in experimentally inoculated ferrets that develop mild respiratory disease, increases the likelihood that virus can spread to the brain via the olfactory nerve as observed in influenza A virus-infected ferrets (36) or in SARS-CoV-2 infected hamsters (17).

Subtle changes were observed in the brains of SARS-CoV-2 inoculated ferrets, despite the lack of evidence for neuroinvasion. These changes included microgliosis and a decrease in astrocytic activation, as well as, an increase in Alzheimer type 2 astrocytes and were present in the acute and post-acute phase. Microglia and astrocytes can be reactive upon direct infection or direct stimuli in their environment such as the presence of viruses or other pathogens in surrounding cells (17,37–42). Another way for increased activation is through indirect stimuli like proinflammatory cytokines caused by a systemic response to for example viral infection (40,41), or a combination of all above. Increased microglial activation upon SARS-CoV-2 infection was observed in patients and experimentally inoculated mice and hamsters, which can be models for severe respiratory disease (7,11,17,23–28). Microglia play a crucial role in maintaining brain homeostasis in health and disease, synapse formation, neuronal proliferation, and brain development. Dysregulation of these processes can impact learning and memory, and deficits have been associated with increased and prolonged microglial activation after infection with SARS-CoV-2, but also with other viruses like influenza A viruses (43–45).

The role of astrocytic activation and Alzheimer type II astrocytes triggered by SARS-CoV-2 infection has been less studied. One study reported an increase in astrocytic activation (28), while another study reported no differences in a hamster model (24). The functional consequences of decreased astrocyte activation are currently unknown, but altered astrocyte physiology can modulate both astrocytic and neuronal glutamate transporter trafficking and activity (46). Thus, a reduction of the reactive astrocyte marker GFAP, meaning that the astrocytes are in a less reactive state, can play a role in dysregulation of glutamate signaling. Altered glutamate signaling is implicated in neuropsychiatric disorders and neurodegenerative diseases, which are both associated with SARS-CoV-2 infection (26,40,43,47,48). Alzheimer type II astrocytes, which we detected in the hindbrain, have recently been detected in the brain of a COVID-19 infected patient. This patient was diagnosed with viral meningoencephalitis with symptoms like altered consciousness and seizures (25). Alzheimer type II astrocytes may be an indicator of hepatic encephalopathy (35,49) but have not been studied extensively. The implications of the presence of Alzheimer type II astrocytes after SARS-CoV-2 infection are currently unknown and need to be studied more.

Together, these data suggest that the ferret serves as a model for self-limiting to mild respiratory disease with limited neuroinvasive potential. Even though the lack of evidence for neuroinvasion, the infection still induced neurovirulence evident by microglial activation, astrocytic deactivation and Alzheimer type II astrocytes. So far, the neuropathogenesis has mainly been studied in animal models for severe disease, while this study highlights the value of this ferret model to study the effects on the brain after mild respiratory disease following intranasal SARS-CoV-2 infection in the acute and post-acute phase.

## Acknowledgements

This work was funded in part by a fellowship from the Netherlands Organization for Scientific Research (VIDI contract 91718308). Ferret experiments were previously published and supported by funding from the National Institutes of Health (AI146980, AI121349, NS091263, AI114736, HHSN272201400008C).

## Author Contributions

**Conceptualization:** Feline F. W. Benavides, Debby van Riel, Lisa Bauer

**Data Curation:** Feline F. W. Benavides, Lonneke Leijten

**Formal analysis:** Feline F. W. Benavides, Edwin J. B. Veldhuis Kroeze, Lonneke Leijten

**Funding Acquisition:** Rory de Vries, Debby van Riel

**Investigation:** Feline F. W. Benavides, Lonneke Leijten, Edwin J. B. Veldhuis Kroeze, Peter van Run

**Methodology:** Katharina S. Schmitz, Rory D. de Vries, Feline F. W. Benavides, Lonneke Leijten

**Project Administration:** Feline F. W. Benavides, Katharina S. Schmitz, Rory D. de Vries, Debby van Riel, Lisa Bauer

**Resources:** Katharina S. Schmitz, Rory D. de Vries

**Software:** Feline F. W. Benavides

**Supervision**: Debby van Riel, Lisa Bauer, Thijs Kuiken

**Validation:** Edwin J.B. Veldhuis Kroeze, Feline F. W. Benavides, Lonneke Leijten

**Visualization:** Feline F. W. Benavides

**Writing – Original Draft Preparation:** Feline F. W. Benavides, Lisa Bauer, Thijs Kuiken, Debby van Riel

**Writing – Review & Editing:** all authors

